# The rates of adult neurogenesis and oligodendrogenesis are linked to cell cycle regulation through p27-dependent gene repression of SOX2

**DOI:** 10.1101/2022.01.25.477676

**Authors:** Ana Domingo-Muelas, Jose Manuel Morante-Redolat, Verónica Moncho-Amor, Antonio Jordán-Pla, Ana Pérez-Villalba, Pau Carrillo-Barberà, Germán Belenguer, Eva Porlan, Martina Kirstein, Oriol Bachs, Sacri R. Ferrón, Robin Lovell-Badge, Isabel Fariñas

**Author notes:** Address for correspondence: Isabel Fariñas, Departamento de Biología Celular, Universidad de Valencia, 46100 Burjassot, Spain. Phones: /3246 (lab). These authors contributed equally to this work (Ana Domingo-Muelas, Jose Manuel Morante-Redolat, Verónica Moncho-Amor). Department of Cell and Developmental Biology, Smilow Center for Translational Research, Perelman School of Medicine, University of Pennsylvania, Philadelphia, PA, USA (A.D-M.), IIS Biodonostia, 48013 Bilbao, Spain (V.M-A.), Max Planck Institute of Molecular Cell Biology and Genetics, 01307 Dresden, Germany (G.B.).

## Abstract

Cell differentiation involves profound changes in global gene expression that often have to occur in coordination with cell cycle exit. Because cyclin-dependent kinase inhibitor p27 reportedly regulates proliferation of neural progenitor cells in the subependymal neurogenic niche of the adult mouse brain, but can also have effects on gene expression, we decided to molecularly analyze its role in adult neurogenesis and oligodendrogenesis. At the cell level, we show that p27 restricts residual cyclin-dependent kinase activity after mitogen withdrawal to antagonize cycling, but it is not essential for cell cycle exit. Contrasting gene expression with chromatin accessibility, we find that p27 is coincidentally necessary to globally repress many genes involved in the transit from multipotentiality to differentiation, including those coding for neural progenitor transcription factors SOX2, OLIG2, and ASCL1. Our data reveal both direct association of p27 with regulatory sequences in the three genes and an additional hierarchical relationship where p27 repression of the *Sox2* gene leads to reduced levels of SOX2-downstream targets *Olig2* and *Ascl1. In vivo*, p27 is also required for the regulation of the proper level of SOX2 necessary for neuroblasts and oligodendroglial progenitor cells to timely exit cell cycle in a lineage-dependent manner.

## Introduction

Mammalian stem and progenitor cell activities are hierarchically organized to produce appropriate numbers of specialized cell types during development and adult tissue renewal. In the subependymal zone (SEZ) of the adult mouse brain, astrocyte-like neural stem cells (NSCs) generate transit-amplifying neural progenitor cells (NPCs) that express the stemness-related transcription factor (TF) SOX2 (Sry-related HMG box 2) and a priming combination of neuron and oligodendroglia fate-specifying bHLH TFs, such as ASCL1 (Achaete-scute homologue 1; also known as MASH1), DLX2 (Distal-Less Homeobox 2) or OLIG2 (Oligodendrocyte Transcription Factor 2) (Gotz et al., 2016; Obernier and Alvarez-Buylla, 2019). These NPCs rapidly divide 3-4 times before they give rise to neuroblasts which themselves divide at least once while migrating to the olfactory bulb (OB) where they exit the cell cycle to fully differentiate as interneurons (Ponti et al, 2013; Obernier and Alvarez-Buylla, 2019). NPCs can also give rise to a small population of oligodendroglial progenitor cells (OPCs) that integrate into the corpus callosum (CC) and differentiate into oligodendrocytes (Menn et al, 2006), as well as to some striatal and CC astrocytes (Sohn et al., 2015). Cell fate decisions along a multistep biological process are dictated and sustained by master TFs, chromatin regulators, and associated networks, but these programs need to act in concert with the regulation of cell cycling and cell cycle exit, a coordination that is still poorly understood (Hardwick et al, 2015).

Two families of cyclin-dependent kinase inhibitors (CKIs) inhibit cell cycle progression by blocking the activity of cyclin/cyclin-dependent kinases (CDK) complexes. Members of the INK4 family p16^Ink4a^, p15^Ink4b^, p18^Ink4c^, and p19^Ink4d^ inhibit early G1 CDKs 4 and 6, whereas the more promiscuous members of the Cip/Kip family p21^Cip1^, p27^kip1^, and p57^Kip2^ can interact with all cyclin-CDK complexes throughout the cell cycle (Besson et al, 2008). Previous analyses have shown that p21 regulates self-renewal of subependymal NSCs (Kippin et al, 2005; Marqués-Torrejón et al, 2013; Porlan et al, 2014) whereas p27 (encoded by the *Cdkn1b* gene) appears to act as a cell cycle inhibitor of adult NPCs (Doetsch et al, 2002; Gil-Perotín et al, 2011). *Cdkn1b* mutant mice (p27KO) are larger in size and exhibit hyperplasia in all organs (Fero et al, 1996; Kiyokawa et al, 1996; Nakayama et al, 1996) suggesting that it plays a role in timing the onset of differentiation in many cell lineages. Reported analyses at the single cell level in the developing CNS have indicated that fetal progenitors without p27 undergo one or two extra rounds of cell division before they stop cycling and differentiate, but the underlying mechanism for such a complex cell behavior has not been elucidated (Durand et al, 1998; Raff, 2007; Defoe et al., 2020). Interestingly, CKI p27 plays roles beyond the control of cell cycle core machinery through cyclinE/CDK2 inhibition (Lim and Kaldis, 2015; Defoe et al., 2020). In line with this, p27 has been shown to promote the differentiation of fetal cortical NPCs through its ability to stabilize TF neurogenin 2 (Nguyen et al, 2006) or to repress *Sox2* gene expression in embryonic stem (ES) and pituitary cells (Li et al, 2012; Moncho-Amor et al, 2021). The non-canonical role of p27 on the transcriptional regulation of specific genes could potentially be a mechanism to timely coordinate cell cycle exit with the implementation of expression programs required to drive differentiation or to repress broad developmental potential (Bachs et al, 2018). However, a comprehensive analysis of p27 actions on gene expression in a somatic cell system has not been performed.

Here, we have analyzed transcriptomic and epigenetic programs modulated by p27 during the transition from proliferation to differentiation in neuronal and oligodendroglial subependymal lineages. We show that p27 participates in timing cell cycle exit through CDK inhibition and, when absent, residual levels of CDK activity allow the cells to engage in extra cell cycles even in the absence of mitogens. Furthermore, using an assay for transposase-accessible chromatin followed by sequencing (ATAC-seq) (Buenrostro et al, 2013) together with RNA deep sequencing (RNA-seq), we characterize global expression changes that require p27 during differentiation. With this approach, we have uncovered a p27-dependent repression of *Sox2, Olig2*, and *Ascl1* genes at the onset of differentiation triggered by mitogen withdrawal. Our data reveal a direct association of p27 with regulatory sequences in these three genes and an additional hierarchical relationship where p27 repression of the *Sox2* gene leads to reduced levels of SOX2-downstream targets *Olig2* and *Ascl1. In vivo* analyses in p27-deficient mice with normal and reduced levels of SOX2 indicate that p27 regulates NPC cycling in a SOX2-independent manner whereas the repressive action of p27 in *Sox2* expression is required for the regulated cell cycle exit of neuroblasts and OPCs. Thus, p27 does not appear to be essential for cell cycle arrest, but it can determine the timing of cell cycle exit through regulation of SOX2 levels in both oligodendrocyte and neuron adult lineages.

## Results

### p27 regulates gene expression programs beyond cell cycle at the onset of differentiation

We first set out to obtain a high-resolution portrait of chromatin structure (ATAC-seq) and gene expression (RNA-seq) at the onset of differentiation in the SEZ neurogenic lineage. Neurogenesis at the murine SEZ is very active with thousands of neuroblasts born every day (Obernier and Alvarez-Buylla, 2019), but cell cycle exit is not synchronous, making it difficult to assess the temporal resolution of cell-specific global changes. Nevertheless, differentiation can be recapitulated *in vitro* and its onset triggered by mitogen withdrawal. To do so, neurospheres grown in EGF and bFGF (mitogenic conditions) are disaggregated and seeded as individual cells on Matrigel and, after 2 days *in vitro* (2 DIV) in a medium containing just bFGF, mitogens are withdrawn (**Figure 1A**). Under these conditions, NSC/NPCs stop dividing after 24 h (2+1 DIV) and initiate a program of differentiation into neurons, oligodendrocytes, and astrocytes (Belenguer et al., 2016) (**Figure 1A**). Comparison of ATAC-seq and RNA-seq data between proliferating NSCs (5-day grown neurospheres) and cells at the onset of differentiation (2+1 DIV) revealed a great number of differentially accessible (DA) promoters and many differentially expressed (DE) genes. DE genes displayed specific patterns of epigenetic changes which led to the majority of promoter regions becoming less accessible (**Figure 1B**). After filtering non-significant changes (FDR>0.05), we defined four gene categories according to their combined differential chromatin-RNA profiles: Closed/Open promoter and UP/DOWN-regulated gene expression, being Closed_UP and Closed_DOWN the ones including most of the differential traits (**Figure 1C**). The less numerous set of Closed_UP genes suggested a dynamic transition from non-accessible chromatin to derepressed expression at cell cycle exit.

**Figure 1.**
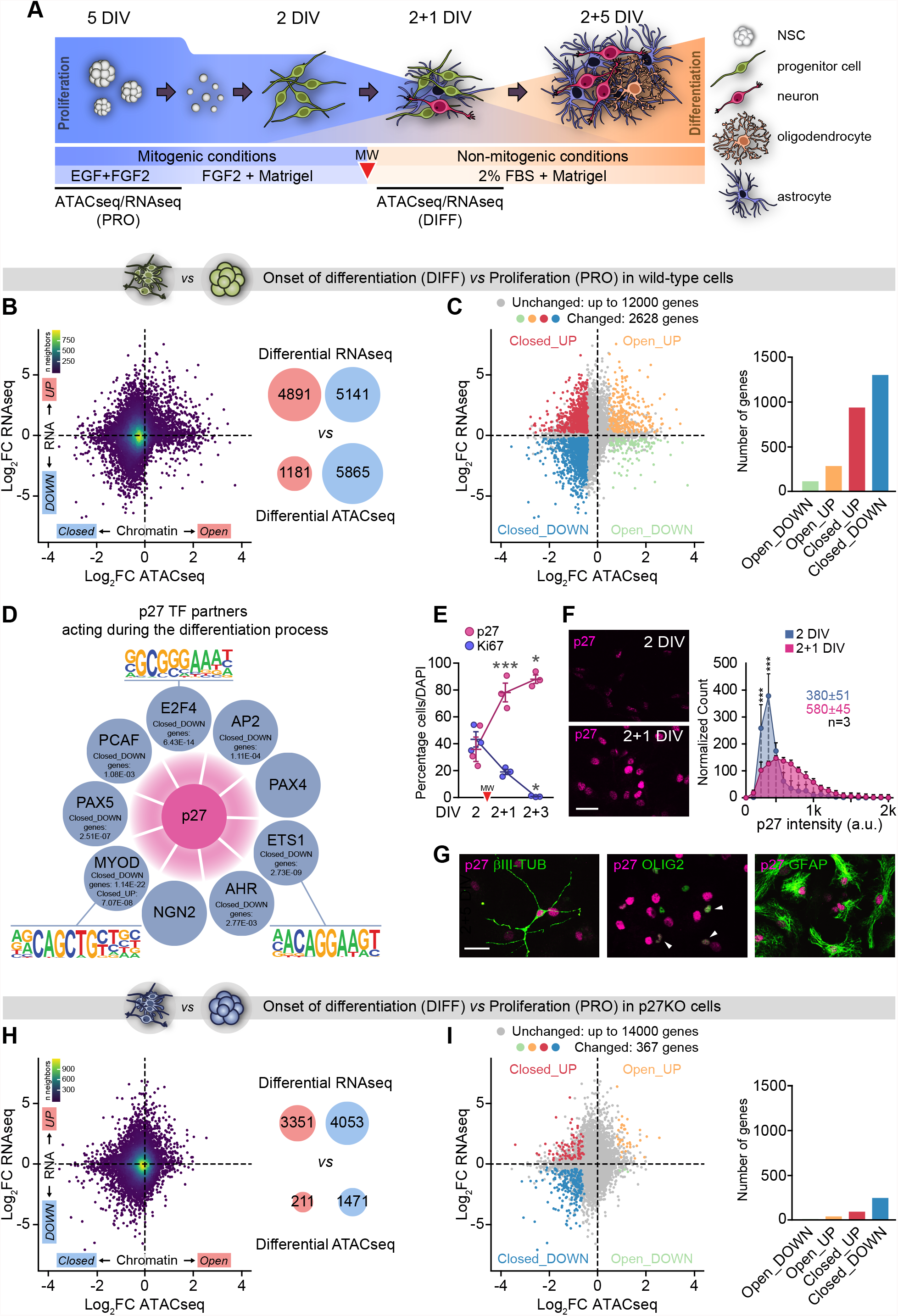
Chromatin and gene expression changes in differentiating NPC cultures that lack p27. **(A)** Schematic drawing of the NSC differentiation protocol. A red arrowhead indicates the critical step of mitogen withdrawal that induces the differentiation process. This symbol will be used equally in the following graphs. MW= mitogen withdrawal, DIV= days *in vitro*. **(B)** Scatterplot of genome-wide chromatin accessibility and mRNA level changes in the transition from proliferation to differentiation in wild type cells. Genes with promoter-associated differential chromatin accessibility are referred to as Open/Closed, whereas genes with differential mRNA expression are referred to as UP/DOWN. Dots are colored according to the density of neighboring dots. Red and blue bubbles indicate the number of genes with statistically significant differential regulation (FDR-controlled p-value < 0.05). **(C)** Scatterplot from B in which genes are coloured according to their integrated differential ATAC-seq and RNA-seq status. The histogram and pie-chart visually summarize the numbers and proportions of genes in the four differential regulation categories: Closed_UP, Closed_DOWN, Open_UP and Open_DOWN **(D)** All known p27 TF partners except two (PAX4, NGN2) were identified by LISA as potential regulatory TFs of genes with dynamic changes in the differentiation process of wild type cells involving reduced chromatin accessibility at promoters. The list of genes to which regulatory TFs were associated is indicated along with their significance values (FDR-controlled p-value). DNA sequence motif logos found by HOMER for three p27 regulatory partners (E2F4, ETS1 and MYOD). **(E)** Quantification of the percentage of p27^+^ (p-value= 0.0002) and Ki67^+^ (p-value=0.0006) cells during differentiation in wild-type cultures. **(F)** Immunocytochemistry showing expression of p27 (magenta) at 2 and 2+1 DIV (left panel). Quantification showing the expression distribution of p27 at 2 and 2+1 DIV. Colored numbers are the median intensity of each time point (right panel). **(G)** Immunocytochemistry (green) for βIII-TUBULIN^+^ neurons, OLIG2^+^ oligodendrocytes and GFAP^+^ astrocytes and for p27 (magenta) at the end of the differentiation protocol (2+5 DIV). **(H) and (I)** Scatterplot of genome-wide chromatin accessibility and mRNA level changes in the transition from proliferation to differentiation in p27KO cells. *p<0.05; ***p<0.001. Scale bars: 30 µm.

We next used the LISA tool (Qin et al, 2020) to infer, through integrative modeling of public chromatin accessibility and ChIP-seq published data, the key putative transcriptional regulators whose combined activity could explain the observed changes. To assemble a list of candidate cis-regulatory regulators for chromatin Closed regions, we used as input for LISA the lists of Closed_UP and Closed_DOWN genes and compared the chromatin landscape models for the target gene sets to background control gene sets. Interestingly, we found several of the top predicted cis-regulators to be known partners of p27 (Bachs et al, 2018) (**Supplementary Table 1**). Based on this, promoter sequence scanning with HOMER revealed consensus sequences for most of the reported p27 TF partners, such as E2F4, MYOD or ETS1, at the promoters of Closed_DOWN genes, suggesting a putative generalized repressive transcriptional role of p27 during differentiation (**Figure 1D**). In line with this, we found that p27 levels sharply increased at 2+1 DIV, when cells stopped proliferation and became Ki67-negative, and remained elevated in differentiated cells (**Figure 1E-G**).

To gain insight into the role of p27, we applied the same ATAC/RNA-seq approach to cultures obtained from p27KO mice (Fero et al, 1996). The number of changes during the differentiation process was remarkably smaller in the absence of p27, specially in ATAC-seq (1,682 *vs* 7,046 DA promoters and 7,404 *vs* 10,032 DE genes; **Figure 1H,I**) resulting in an 85% reduction in the number of genes with changes in either expression or accessibility (367 in p27KO *vs* 2,628 in WT). Intersection of the lists of genes with promoter/mRNA changes in both genotypes indicated that 88% of the changes in DE genes showing Closed chromatin in the WT process were lost in the absence of p27 (**Figure 2A**). Since our results indicated a generalized chromatin repression after mitogen withdrawal that appeared largely lost in p27KO cells, we performed functional enrichment analysis for genes in the Closed category during differentiation. Gene Ontology (GO) analysis identified categories of genes related to “cell cycle control” in the Closed_DOWN group whereas most genes in the Closed_UP group were related to “neuronal differentiation/neurogenesis”. Importantly, both categories were affected by the loss of p27 (**Figure 2B**). Therefore, the generalized loss of epigenetic and expression changes and the identity of the affected genes suggested that p27 could act as an important factor in establishing the appropriate chromatin and regulatory landscapes for proliferating neural cells to differentiate, acting on cell cycle and differentiation genes.

**Figure 2.**
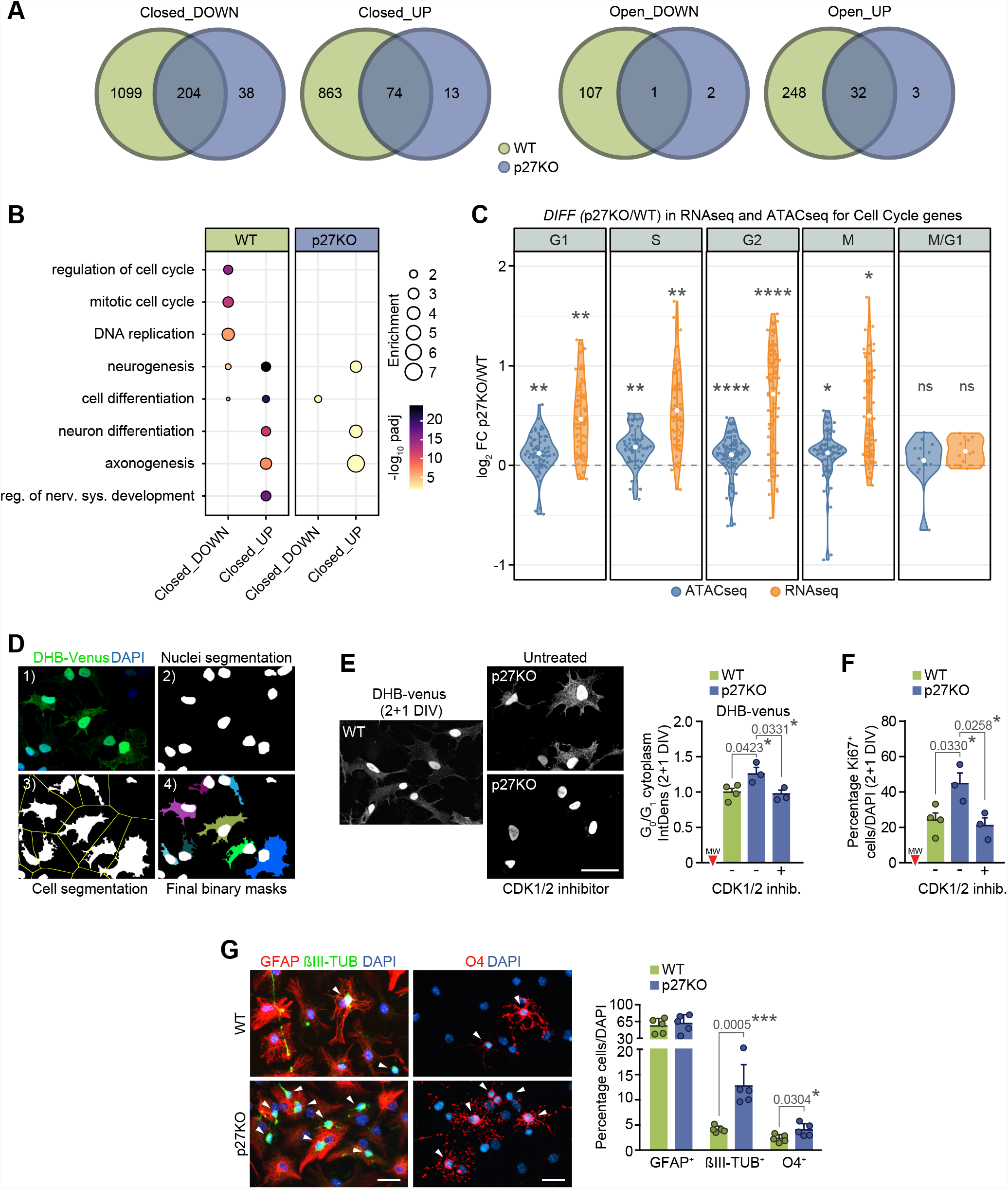
Cell cycling and cell fate decisions in adult NPC cultures are regulated by p27. **(A)** Venn diagrams comparing the number of genes with simultaneous changes in chromatin and expression in wild type and p27KO cells. **(B)** Dot plot representation of the functional enrichment analysis of the genes in the overlapping regions shown in A. **(C)** Violin plots showing the changes between p27KO and wild type cells in promoter accessibility and expression of genes involved in the different phases of the cell cycle. **(D)** Main steps during DHB-Venus bioimage analysis workflow (see Material and Methods for detailed explanation). **(E)** DHB-venus signal (white) in WT and p27KO cultures at 2+1 DIV after CDK1/2 inhibitor treatment. Comparison of cytosolic grey integrated density (IntDens) of DHB-Venus fluorescence of cells in G_0_/G_1_ measured by Bioimage Analysis. Wild-type, and p27-deficient cultures, either untreated or treated with 1 µM CDK1/2 inhibitor were imaged and scored during differentiation at 2+1 DIV. Data is represented as fold change relative to wild-type. **(F)** Quantification of the percentage of Ki67^+^ cells at 2+1 DIV in WT, p27KO and mutant cultures treated with 1 µM CDK1/2 inhibitor. **(G)** Immunocytochemistry (left) and quantification (right) of βIII-TUBULIN^+^ neurons (green), GFAP^+^ astrocytes (red) and O4^+^ oligodendrocytes at 2+5 DIV (left) in WT and p27KO cultures. Arrowheads point out positive cells. DAPI was used to counterstain nuclei. Graphs represent mean and all error bars show s.e.m. The number of independent biological samples used is indicated as dots in the graphs. *p<0.05; **p<0.01; ***p<0.001. Scale bars: 30 µm.

The CKI p27 plays a canonical inhibitory role in cell cycle regulation. Therefore, as a first approach to understand the specific gene expression programs that failed to trigger in the absence of p27, we analyzed RNA and ATAC data separately for genes characteristic of each cell cycle phase (Li et al, 2019) by calculating the fold change between p27KO and WT. The data indicated persistent expression of cell cycle genes in the absence of p27 after mitogen withdrawal (**Figure 2C**), in accordance with higher proportions of EdU-incorporating cells (9.1 ± 2.0, n = 5, *vs* a wild-type value of 2.2 ± 0.4, n = 6, p-value = 0.0015). Cell cycle phase-dependent changes in the transcription of genes are regulated by CDK-dependent phosphorylation of retinoblastoma protein, RNA polymerase II and some specific TFs (Bachs et al, 2018) and, therefore, observed changes in cell cycle gene expression in the absence of p27 could be CDK-dependent as cell cycling itself. Indeed, the decision to re-enter the cell cycle is reportedly controlled by the residual level of CDK2 activity at the point of mitotic exit (Overton et al, 2014; Spencer et al, 2013). We, therefore, took advantage of the CDK2 reporter CSII-EF-DHB-mVenus, composed of the C-terminal CDK2-phosphorylation domain of the human DNA helicase B fused to the yellow fluorescent protein mVenus (Spencer et al, 2013). The reporter is primarily found in the nucleus during mitosis and in newly generated cells, but it translocates to the cytoplasm upon phosphorylation by CDK2 during G1/S transition. Quantitative image analysis of the mVenus cytoplasm/nucleus (C/N) ratio in cells transduced with the reporter (**Figure 2D**) indicated that p27-deficient cells that were in G0/G1 (C/N <0,95) displayed higher cytosolic fluorescence than controls at 2+1 DIV, reflecting increased CDK2 activity (**Figure 2E**). In agreement with these data, cytosolic Venus fluorescence and the proportion of Ki67^+^ cells in p27KO cultures deprived of mitogens could be reduced to wild-type levels by treatment with a pharmacological CDK1/2 inhibitor (**Figure 2E,F**). This result indicated that p27-dependent inhibition of CDK2 at cell cycle exit restricts cell cycle progression and suggested that effects of p27 in cell cycle gene expression may likely be also dependent on CDK activity.

Our results indicated that, in the absence of p27, residual levels of CDK2 activity endow some cells with the potential to commit to another cell cycle even in the absence of mitogens. Despite the CDK-dependent inhibitory effect of p27 on cycling, the CKI does not appear to be essential for cell cycle arrest of adult NPCs as cells fully stop proliferation and fully differentiate by 2+5 DIV in the absence of p27 (**Figure 2G**), in line with reports in other cell types (Durand et al., 1998). Interestingly, although p27-deficient NPCs were overall capable of engaging extra cell divisions, mutant cultures produced higher proportions of neurons and oligodendrocytes, but not astrocytes (**Figure 2G**). This population-specific effect, together with the detected changes in gene expression and accessibility in the Closed category related to GO terms of “neuronal differentiation/neurogenesis”, suggested that the role of p27 could go beyond the cell cycle to affect differentiation through specific actions on gene expression.

### p27 negatively regulates a SOX2-Olig2/Ascl1 axis at cell cycle exit

To address the possibility that p27 was establishing a timer mechanism for differentiation through the modulation of a transcriptional landscape in NPCs, we next sought to identify the network of differentiation-driving transcriptional regulators acting under the control of p27. We analyzed the lists of differentiation-associated Closed genes of both wild-type and p27KO cells (**Figure 3A**) with the LISA tool in order to find TFs already reported in the literature to bind to those genes (**Figure 3B**) and with the HOMER tool to infer the potential regulators by detecting the presence of binding sites in the promoters of the genes in the lists (**Figure 3C**). Comparative analysis of the LISA results in wild-type *vs* p27KO revealed that the three most relevant TFs among the top 25 were ASCL1, OLIG2 and SOX2 and that their influence was more pronounced in the mutants (**Figure 3B; Supplementary Table 2**). This result was corroborated by HOMER which identified a set of 49 TFs likely controlling the differentiation in both genotypes but also a group of 21 that were more prominently acting when p27 was absent. The interaction network of these p27KO-specific TFs placed ASCL1, OLIG2 and SOX2 at the core of the network (**Figure 3C**). Taken together, the data suggested that p27 is required at the onset of differentiation to restrict the presence of SOX2, OLIG2 and ASCL1 in a subset of repressed genes.

**Figure 3.**
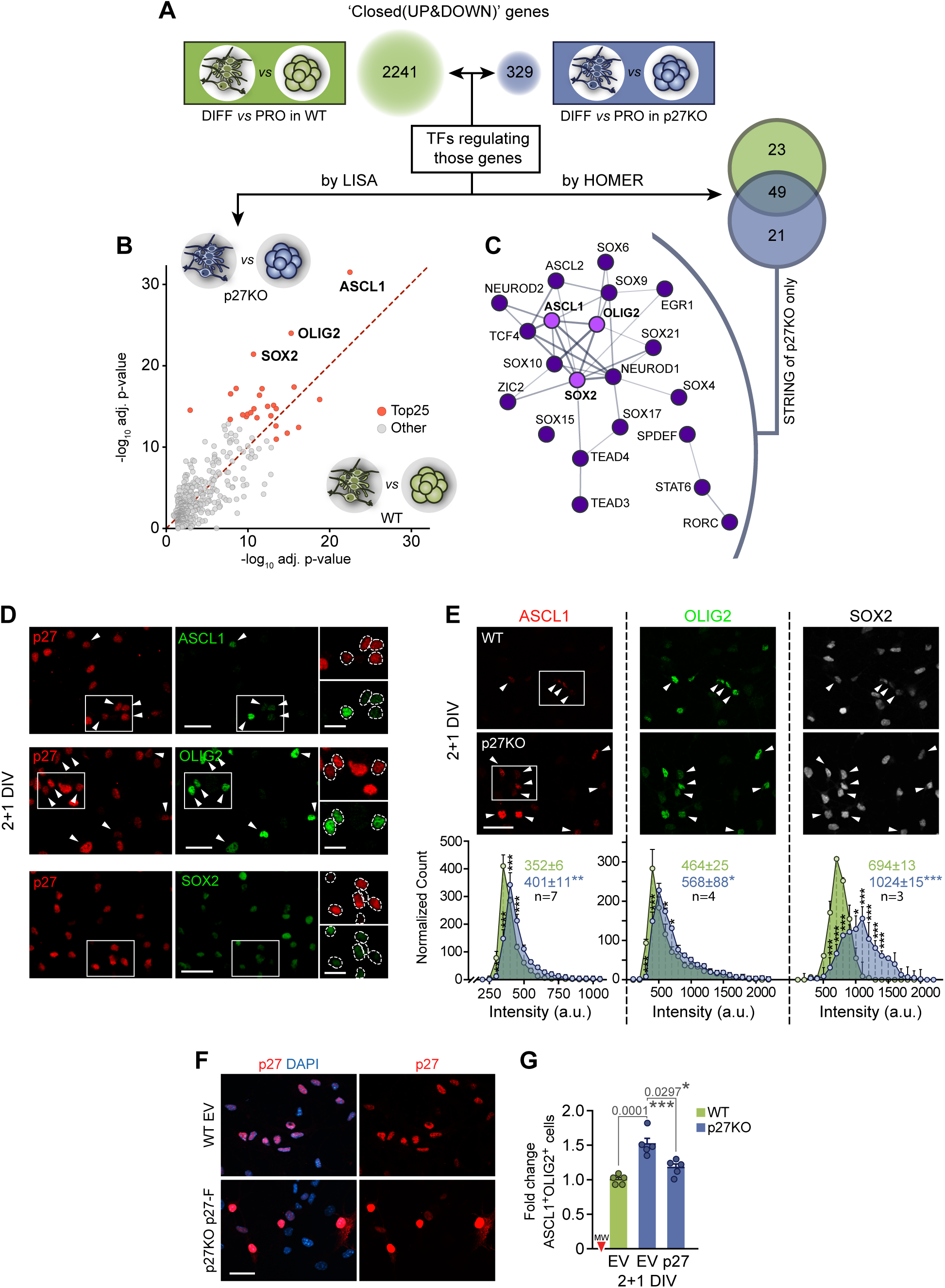
Deregulation of SOX2, ASCL1 and OLIG2 TFs in p27KO cultures at the onset of differentiation. **(A)** Schematic drawing of the genotypes compared with the integrative genomic analysis of RNA-seq and ATAC-seq datasets: differentiating vs proliferating wild type cells (green), or p27KO cells (blue) are shown. Numbers in bubbles indicate, for each of the comparisons, the amount of differentially expressed genes (UP and DOWN-regulated) with decreased promoter chromatin accessibility. **(B)** Both gene lists were comparatively analyzed with LISA to search for putative regulatory TFs which might be involved in their regulation. The scatterplot shows the statistical significance with each of the potential regulatory TFs associated with the genes in each of the lists. **(C)** The promoter sequences of genes in both lists were scanned with HOMER to look for TF binding site motifs. The Venn diagram shows the overlap between the statistically significant (FDR < 0.05) TF binding sites found for each comparison. The interaction network shows the relationships between the 21 TFs with binding sites at the promoters of differentially regulated genes exclusively in the transition between proliferation and differentiation in p27KO cells. **(D)** Immunocytochemistry for p27 (red) in combination with ASCL1, OLIG2 or SOX2 (green) in 2+1DIV NPs. **(E)** Immunocytochemistry for the transcription factors ASCL1 (red), OLIG2 (green) and SOX2 (white) in 2+1DIV WT and p27KO NPCs (top panels). Quantification showing the expression distribution of each transcription factor as a result of p27 deficiency. Colored numbers are the median intensity of each genotype (bottom panels). **(F)** Immunostaining for p27 (red) in wild-type cultures transfected with an empty vector (EV) and p27KO cultures overexpressing a full-length p27 construct (p27-Flag) (2+1 DIV). **(G)** Quantification of the ASCL1^+^OLIG2^+^ population that expresses SOX2 in WT and p27KO cultures at 2+1 DIV. knock-out cultures after reintroduction of a full length p27-construct are also quantified. An empty vector (EV) was used as a negative control. Data is represented as fold change relative to wild-type. The number of independent biological samples used is indicated as dots in the graphs. Graphs represent mean and all error bars show s.e.m. Exact p-values are indicated in the graphs and legend, being *p<0.05; **p<0.01; ***p<0.001. Scale bars: 30 µm (inserts in D: 15 µm).

Identifying SOX2, OLIG2 and ASCL1 as the most prominent TFs potentially involved in the altered differentiation of p27KO cells led us to investigate whether p27 could be acting as a regulator of the corresponding genes. SOX2 is expressed by NSCs and NPCs (Reiprich & Wegner, 2015; Sarkar & Hochedlinger, 2013). ASCL1 is expressed by activated NSCs and by NPCs for both oligodendrocytes and neurons, whereas the presence of OLIG2 in a few NPCs renders them oligodendrogenic (Parras et al, 2004; Hack et al, 2005; Colak et al., 2008). Around 40% of all cells in a proliferating culture are SOX2^+^OLIG2^+^ASCL1^+^ but this percentage becomes reduced at 2+1 DIV to around 10% (41.0 ± 2.6% at 2 DIV *vs* 12.2 ± 0.7% at 2+1 DIV, n = 4, p-value = 0.002), in inverse correlation with the increasing levels of p27 (**Figure 1E-F; 3D**). In agreement with a repressive action of p27, we scored a significantly higher proportion of cells with increased levels of ASCL1, OLIG2 and SOX2 in differentiating p27KO cells that could be restored to wild-type levels by transduction of a p27 cDNA (**Figure 3E-G**).

We next set up to evaluate direct physical interaction of p27 with regulatory regions in their coding genes under proliferative and differentiative conditions. Highly conserved regulatory regions in the *Sox2* gene include a core promoter (Wiebe et al, 2000) and a number of enhancers organized in Sox2-regulatory regions (SRR), where SRR1 contains enhancers (N) 2 and 3 and SRR2 contains N1, 4, and 5 (Miyagi et al, 2004; Zappone et al, 2000) and is active in the adult SEZ (Miyagi et al, 2006). As for *Olig2* and *Ascl1*, the *in silico* analysis of the 2kb sequence upstream of their transcription initiation site combining Genomatix and EPD tools revealed the presence of several putative binding sites for E2F4 and ETS1 TFs, some of them clustered in the proximal promoter (−1kb) (**Figure 4A**). Therefore, we performed chromatin immunoprecipitation (ChIP) with anti-p27 antibodies in wild-type cultures to detect the binding of p27 to several of these regulatory regions (Sox2: [-460, -345], Sox2 SRR2: [+4042, +4133], Olig2: [-637, -546] and [-147, -58], Ascl1: [-906, -784] and [-523, -432]). In proliferative cells we could detect binding of p27 to the *Sox2* proximal promoter and the SRR2 enhancer and to the proximal promoters of *Olig2* and *Ascl1* (ratio p27-IP over NRA control ranging between 1.82 and 3.70). At the onset of differentiation, when p27 levels increased abruptly, the presence of p27 in the *Sox2* proximal promoter declined, whereas binding to the SRR2 enhancer as well as to all the tested regulatory regions in *Olig2* and *Ascl1* promoters was strengthened 2-3 fold (**Figure 4A**). The results on the *Sox2* promoter and SRR2 enhancer were confirmed by means of specific luciferase reporters: the activities of Sox2prom[-1907,+6]-luc and SRR2[+3641/+4023]-luc) (Tomioka et al, 2002) constructs were higher in proliferating mutant cells, dropped or increased upon differentiation respectively, and could both be repressed by reintroduction of p27 (**Figure 4B**).

**Figure 4.**
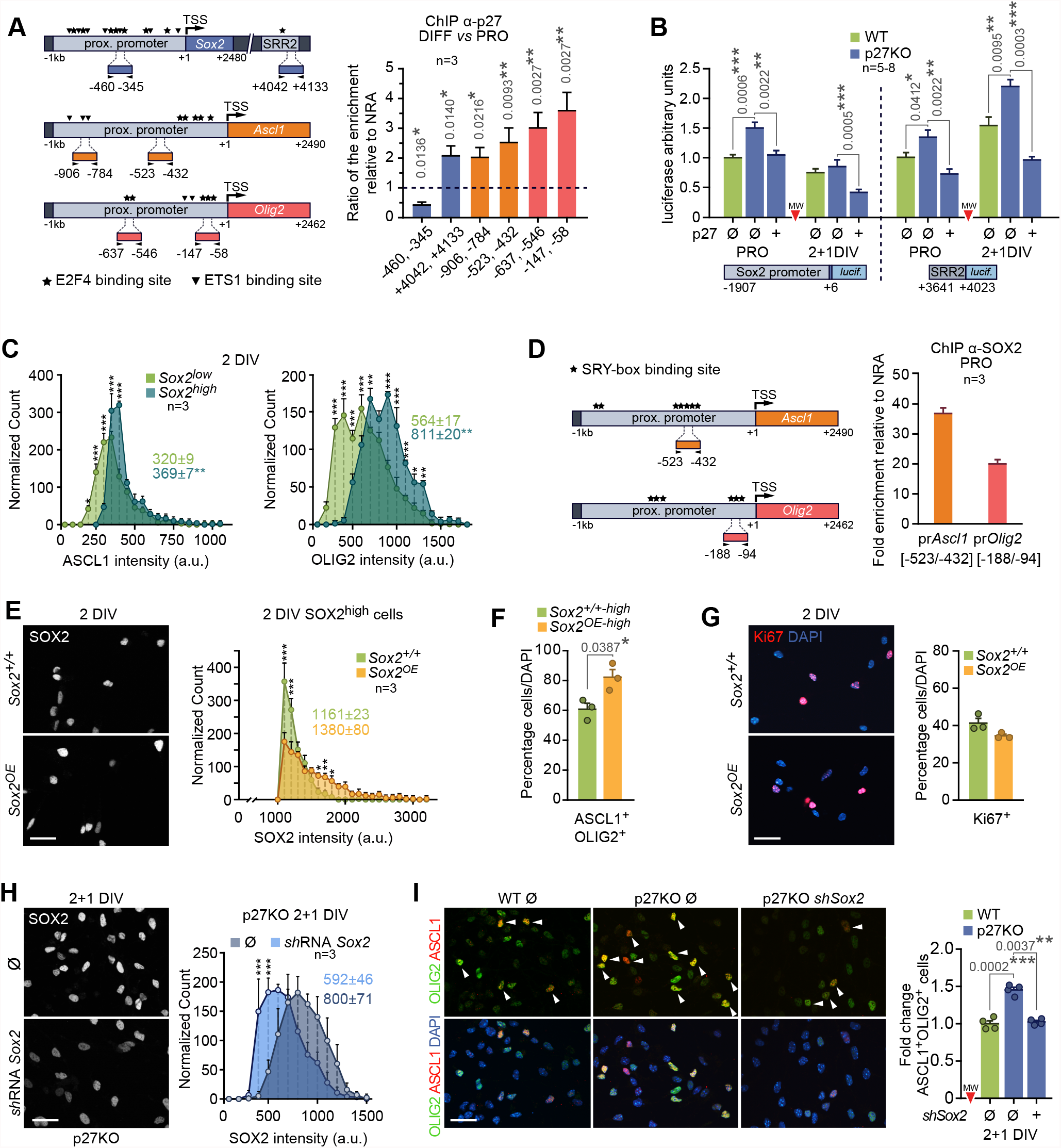
Direct and indirect regulation of Sox2, Ascl1 and Olig2 genes by p27. **(A)** Schematic drawing of *Sox2, Ascl1* and *Olig2* genes with the amplicons used in the p27ChIP protocol. TSS: Transcription Start Site. Stars represent putative E2F4 binding sites whereas invered triangles represent putative ETS1 binding sites (left panel). ChIP assay of p27 in differentiative (DIFF, 2+1 DIV) relative to proliferative (PRO, spheres) conditions. Binding to each regulatory region is represented as the ratio of enrichment normalized to a non-related antibody (NRA) (right panel). **(B)** Luciferase reporter assay in PRO and DIFF conditions in wild-type, p27KO cultures, and p27KO cells transfected with a full-length p27 construct. Transcription activity for both the proximal promoter (*prSox2*[-1907/+6]) and the *SRR2* enhancer (*SRR2*[+3641/+4023]) is represented as arbitrary units relative to wild-type during proliferation. **(C)** Frequency histograms showing ASCL1 and OLIG2 levels among SOX2^low^ and SOX2^high^ populations at 2 DIV of differentiation. Colored numbers are the median intensity of each population. **(D)** Schematic drawing of *Ascl1* and *Olig2* genes with the amplicons used in the SOX2 ChIP protocol. Stars represent putative binding sites for the SOX family (SRY-box bs) (left panel). ChIP assay of SOX2 during proliferation. Binding to the *Ascl1* promoter (*prAscl1*[-522/-432]) and the *Olig2* promoter (*prOlig2*[-188/-94]) is represented as fold enrichment relative to a non-related antibody (NRA) (right panel). **(E)** Immunocytochemistry showing SOX2 expression (white) after Sox2 overexpression at 2 DIV (left panel). Histogram showing SOX2 levels in the 30% brightest cells (SOX2^high^) of wild-type and overexpressing (*Sox2*^*OE*^) cultures at 2 DIV. Colored numbers indicate the median intensity (right panel). **(F)** Quantification of the percentage of ASCL1^+^OLIG2^+^ cells among the SOX2^high^ population during 2 DIV of differentiation of wild-type (*Sox2*^*+/+-high*^) and SOX2 overexpressing cultures (*Sox2*^*OE-high*^). **(G)** Representative images and quantification of the percentage of Ki67^+^ progenitors at 2 DIV after SOX2 overexpression (p-value = 0.1552). **(H)** Immunocytochemistry showing SOX2 expression (white) (left panel) and the corresponding quantification (right panel) in p27KO cultures after *Sox2* downregulation by *sh*RNA*Sox2* at 2+1 DIV. Colored numbers indicate the median intensity (p-value=0.0604). **(I)** Representative images (left) and quantification (right) of the ASCL1^+^OLIG2^+^ population at 2+1 DIV after transfection with a *sh*RNA*Sox2* in p27-deficient cells. An empty vector (Ø) was used as a negative control in WT and p27KO cultures. Data is represented as fold change relative to wild-type. Graphs represent mean and all error bars show s.e.m. The number of independent biological samples used is indicated as dots in the graphs. Exact p-values are indicated in the graphs and legend, being *p<0.05; **p<0.01; ***p<0.001. Scale bars: 30 µm

Interestingly, the levels of OLIG2 and ASCL1 quantitatively correlated with those of SOX2 at the single cell level (**Figure 4C**), suggesting the possibility that p27 could also be exerting its transcriptional repression effect on *Olig2* and *Ascl1* indirectly through its control of *Sox2*. Indeed, when we performed ChIP with anti-SOX2 antibodies, we could detect strong binding to the proximal promoters of both TFs (Olig2 [-188, -94], Ascl1 [-522, -432]) (**Figure 4D**). Consistently, overexpression of Sox2 by viral delivery of a Cre-recombinase to cultures obtained from R26-loxP-STOP-loxP-Sox2-GFP mice (Lu et al, 2010) resulted in increased proportions of ASCL1^+^OLIG2^+^ cells without effects on proliferation (**Figure 4E-G**). The hierarchical relationship of SOX2 over *Olig2* and *Ascl1* suggested the possibility that the increase in the proportion of ASCL1^+^OLIG2^+^ NPCs resulting from a lack of p27 could be rescued by lowering SOX2 levels. Indeed, reduction of SOX2 protein levels, following nucleofection of a specific *Sox2* shRNA in p27-deficient cultures, rescued the proportion of OLIG2^+^ASCL1^+^ NPCs as efficiently as reintroducing p27 (**Figure 4H,I**). Our data together indicated that p27 can potentially act as a repressive regulator of the *Sox2, Olig2*, and *Ascl1* genes through physical binding but that, functionally, its direct repressive action on the *Sox2* gene is sufficient to reduce the downstream expression of *Ascl1* and *Olig2*.

### Regulation of Sox2 by p27 is required for timely cell cycle exit in adult neurogenesis and oligodendrogenesis in a cell context-dependent manner

Next, we decided to test the uncovered regulation of these three TFs by p27 and its functional consequences in the intact SEZ. In postnatal mice, p27 reportedly exhibits a generalized expression in migrating neuroblasts and in young OB neurons with a caudal^low^-rostral^high^ gradient (Li et al, 2009), but distribution of the protein in the adult was so far unknown. Analysis with antibodies specific to p27 in young adult (2 month-old) male and female mice indicated that the protein is barely detectable in adult GFAP^+^ NSCs and shows low and variable levels in ASCL1^+^ NPCs, while a strong signal is detected in DCX^+^ cells (**Figure 5A,B**), suggesting that p27 levels increase with lineage progression and is maximal in neuroblasts.

**Figure 5.**
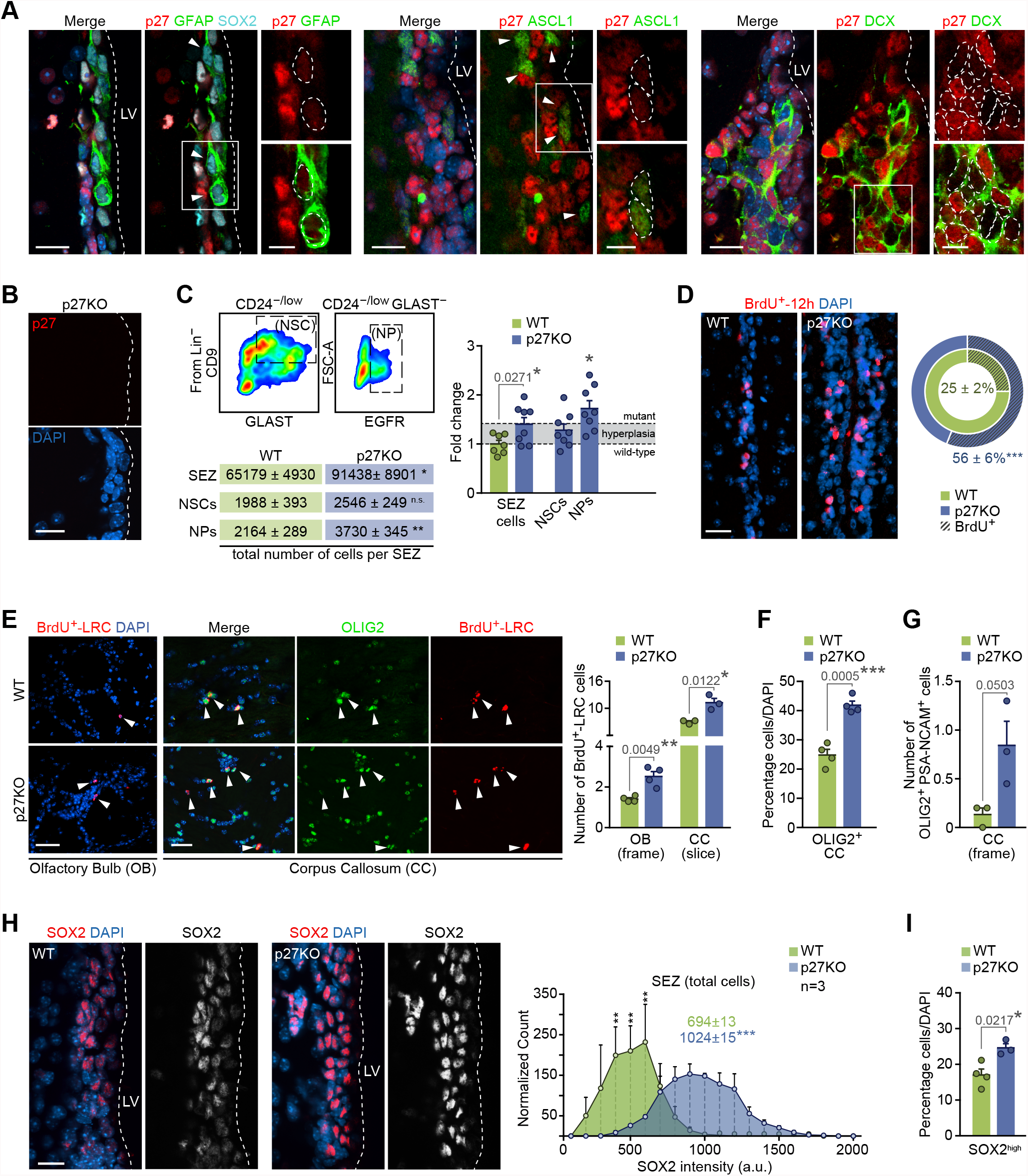
p27 determines the rate of NPC-sustained neurogenesis and oligodendrogenesis in the adult SEZ. **(A)** Immunohistochemistry showing expression of p27 (red) in GFAP^+^(green)/SOX2^+^ (cyan) NSCs, ASCL1^+^ (green) NPs and DCX^+^ (green) NBs in the SEZ of wild-type mice. **(B)** Immunohistochemistry for p27 (red) in the SEZ of p27KO mice. **(C)** Representative FACS plots showing GLAST and CD9 staining of Lin^−^ CD24^−/low^ fraction where NSCs are identified by CD9^high^ levels in the GLAST^+^ cells while the Lin^−^ CD24^−/low^ GLAST^−^ region contains EGFR^+^ NPs (top left panel). Quantification of total number of NSCs (p-value=0.2400) and NPs (p-value=0.0046) analyzed by FACS in the SEZ of WT and p27KO mice (bottom left panel). Analysis of the total number of SEZ cells and NSCs and NPs populations by flow cytometry represented as fold change relative to wild-type (NSCs: n.s., p-value = 0.5585; NPs: p-value=0.0432). Dashed line represents the observed increment in p27KO number of cells owing to hyperplasia (right panel). **(D)** Immunohistochemistry for BrdU (red) within the SEZ of WT and p27KO mice. Quantification of the percentage of cells that incorporated BrdU in the 12h prior to euthanasia (BrdU^+^-12h) in each genotype. **(E)** Immunohistochemistry showing the staining of BrdU-LRC (red) in the glomerular layer of the olfactory bulb (OB), and of BrdU-LRC (red) and OLIG2 (green) in the *corpus callosum* (CC) of WT and p27KO mice. Arrowheads indicate double positive cells (left panel). Quantification of the number of BrdU-LRC cells in the OB and CC of wild-type and p27-deficient mice (right panel). **(F)** Quantification of the percentage of OLIG2^+^ oligodendrocytes in the CC of WT and p27KO adult brains. **(G)** Quantification of the number of PSA-NCAM^+^OLIG2^+^ cells per frame in the CC of WT and p27KO mice. **(H)** Immunohistochemistry showing the expression of SOX2 (red and white) in the SEZ of wild-type and p27 deficient mice (left panel). Histogram showing quantification of SOX2 intensity. Colored numbers are the median intensity of each population (right panel). **(I)** Quantification of the percentage of cells that express high levels of SOX2 in the SEZ of wild-type and p27 deficient mice. DAPI was used to counterstain nuclei. LV: lateral ventricle. Graphs represent mean and all error bars show s.e.m. The number of independent biological samples used is indicated as dots in the graphs. Exact p-values are indicated in the graphs and legend, being *p<0.05; **p<0.01; ***p<0.001. Scale bars: A-B, H, 20 µm (inserts 10 µm) ; D-E, 30 µm

Adult *Cdkn1b* mutant mice have larger bodies (Fero et al, 1996) and we could also observe enlarged SEZs (volume, in mm^3^ × 10^6^ ± s.e.m.: 21.3 ± 1.8 *vs* a wild-type value of 15.3 ± 0.9, n = 3, p-value = 0.0411) and OBs (6.0 ± 0.1 *vs* a wild-type value of 4.7 ± 0.1, n = 3, p-value = 0.0007). In line with this, SEZ homogenates showed a 40% increase in cell yield (**Figure 5C**); however, when we scored specific cell populations by flow cytometry (Belenguer et al, 2021a,b), we found that the increase in NSCs (CD24^−/low^GLAST^+^CD9^high^ cells) was within the range of the generalized hyperplasia of the tissue, whereas NPCs (GLAST^−^CD24^−/low^EGFR^+^ cells) were significantly overrepresented with an increase of 70% in *Cdkn1b* mutant mice (**Figure 5C**). In agreement with this, 2-month old mice intraperitoneally injected with 7 pulses of BrdU, one every two hours, during the 12h period preceding euthanasia, evidenced that overall BrdU-incorporation rate doubled in p27KO mice (**Figure 5D**), whereas GFAP^+^ cells exhibited a normal BrdU-labelling rate (percentage of GFAP^+^ that were BrdU^+^ ± s.e.m.: 8.9 ± 2.0 vs a wild-type value of 10.2 ± 1.6, n = 3, p-value = 0.5980). The data confirmed that p27 is a specific regulator of NPC cycling, as previously reported (Doetsch et al, 2002; Gil-Perotin et al, 2011). Regarding cell progeny output, we found higher proportions of BrdU^+^ label-retaining cells (LRCs) in the OB glomerular layer and in the CC of mice injected with 7 pulses of BrdU four weeks before euthanasia (**Figure 5E**). The increase in the density of LRCs in the CC correlated with more OLIG2^+^ cells that were also positive for PSA-NCAM (characteristic of oligodendroglia generated in the SEZ; Menn et al, 2006) (**Figure 5F,G**). Therefore, p27 limits both neurogenesis and oligodendrogenesis in the adult brain.

Increased production of neurons and oligodendrocytes in the absence of p27 could be the result of an enlarged population of proliferating NPCs in the SEZ. However, our data indicated that p27 negatively regulates the levels of TFs SOX2, ASCL1 and OLIG2, largely by transcriptional repression of the *Sox2* gene. Analysis of the SOX2 protein by quantitative image analysis in immunostained SEZ sections revealed higher levels *per* cell and increased proportions of SOX2^high^ cells also *in vivo* (**Figure 5H,I**). In contrast to the culture situation, OLIG2 is restricted to a minority (around 2%) of ASCL1^+^SOX2^+^ NPCs that behave as OPCs and specifically generate oligodendrocytes *in viv*o (Parras et al, 2004; Hack et al, 2005; Colak et al., 2008). Analyses of these OLIG2^+^ASCL1^+^ cells in p27KO mice revealed increased levels of SOX2 per cell (**Figure 6A**) and higher proportions of OPCs that, furthermore, proliferated more actively as shown by increased BrdU incorporation rate after a 1 h-chase (**Figure 6B-D**). As with regards to the neuronal lineage, doublecortin (DCX)^+^ early neuroblasts generated from NPCs can proliferate once or twice as they retain EGFR and are sensitive to mitogens before they exit cell cycle as late non-dividing EGFR^-^ neuroblasts that constitute, by far, the largest population in the SEZ neurogenic niche (Ponti et al, 2013; Belenguer et al, 2021). We found that DCX^+^ neuroblasts in wild-type mice had very low or undetectable levels of SOX2 and ASCL1 (Baser et al, 2019; Ferri et al, 2004) together with high levels of p27 (**Figure 5A,B**), whereas SOX2 and ASCL1 levels remained abnormally high in a fraction of p27KO neuroblasts (**Figure 6E,F**). Although the absence of p27 also resulted in increased proportions of neuroblasts in the SEZ, it did not alter their proliferation rate, as determined by a 1-h chase after a single injection of BrdU (**Figure 6G,H**). The results together indicated that, *in vivo*: 1) p27 regulates the cycling of NPCs, including those of oligodendrocytes, but not of neuroblasts, and 2) p27 restricts the levels of SOX2 in both OPCs and neuroblasts.

**Figure 6.**
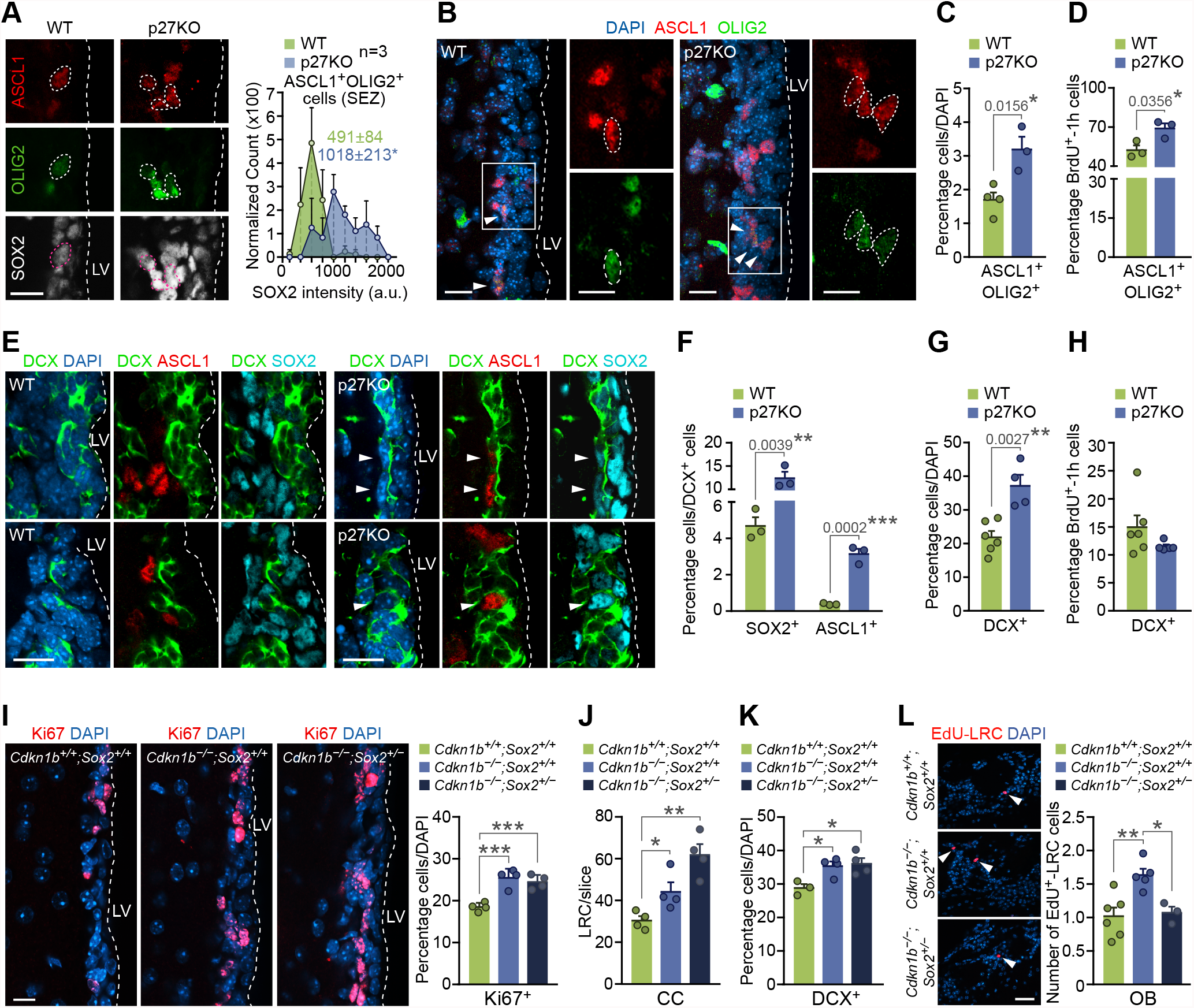
p27-dependent levels of SOX2 regulates timely cell cycle exit in adult neurogenesis and oligodendrogenesis. **(A)** Representative immunohistochemistry images for ASCL1 (red) and OLIG2 (green) and SOX2 (white) in the SEZ of WT and p27KO mice. Triple positive cells have been delineated (left). Quantification of SOX2 expression intensity in ASCL1^+^OLIG2^+^ cells in the SEZ of WT and p27KO mice (right). **(B)** Representative immunohistochemistry images for ASCL1 (red) and OLIG2 (green) in the SEZ of WT and p27KO mice. Arrowheads indicate double positive cells **(C)** Quantification of the percentage of ASCL1^+^OLIG2^+^ cells in the SEZ of adult wild-type and p27-deficient mice. **(D)** Quantification of the percentage of proliferating cells (BrdU-1h^+^) among the ASCL1^+^OLIG2^+^ population as a result of p27 deficiency. **(E)** Immunohistochemistry showing expression of ASCL1 (red), DCX (green) and SOX2 (cyan) in the SEZ of wild-type and p27 deficient mice. Arrowheads indicate triple-positive cells. **(F)** Quantification of the percentage of DCX^+^ neuroblasts that are positive for SOX2 or ASCL1 in the SEZ of wild-type and p27 knock-out mice. **(G)** Quantification of the percentage of DCX^+^ neuroblasts produced in the adult SEZ. **(H)** Quantification of the percentage of proliferating cells (BrdU-1h^+^) among the DCX^+^ p27-mutant population. **(I)** Immunohistochemistry for Ki67 (red) in the adult SEZ of wild-type, p27 knock-out (*Cdkn1b*^*–/–*^*;Sox2*^*+/+*^) and p27-deficient, *Sox2* heterozygous (*Cdkn1b*^*–/–*^*;Sox2*^*+/–*^) mice (left panel). Quantification of the percentage of Ki67^+^ cells (ANOVA p-value=0.0004) (right panel). *(J)* Quantification of the number of EdU-LRC cells in the CC of *Cdkn1b*^*+/+*^*;Sox2*^*+/+*^, *Cdkn1b*^*–/–*^*;Sox2*^*+/+*^ and *Cdkn1b*^*–/–*^*;Sox2*^*+/–*^ mice. **(K)** Quantification of the percentage of DCX^+^ cells in the SEZ of *Cdkn1b*^*+/+*^*;Sox2*^*+/+*^, *Cdkn1b*^*–/–*^*;Sox2*^*+/+*^ and *Cdkn1b*^*–/–*^ *;Sox2*^*+/–*^ mice. **(L)** Immunohistochemistry and quantification of the number of BrdU-LRC (red) reaching the glomerular layer of the OB of *Cdkn1b*^*+/+*^*;Sox2*^*+/+*^, *Cdkn1b*^*–/–*^*;Sox2*^*+/+*^ and *Cdkn1b*^*–/–*^ *;Sox2*^*+/–*^ mice (ANOVA p-value= 0.0051). DAPI was used to counterstain nuclei. LV: lateral ventricle. Graphs represent mean and all error bars show s.e.m. The number of independent biological samples used is indicated as dots in the graphs. Exact p-values are indicated in the graphs and legend, being *p<0.05; **p<0.01; ***p<0.001. Scale bars: A, E, I, 20 µm (inserts 10 µm); L, 30 µm.

Because of the dual action of p27, we next decided to test whether increased levels of SOX2 played a role in the *p27KO* phenotype by analyzing *p27KO*;*Sox2* heterozygous (Avilion et al, 2003) compound double mutant mice. Overall, heterozygous levels of SOX2 did not reduce the increased proportion of Ki67^+^ proliferating cells resulting from the deletion of p27, indicating that cell cycle regulation in NPCs is SOX2-independent (**Figure 6I**). SOX2 effects in neural cells are dosage-sensitive, but also context dependent (Zhang et al., 2018) and, therefore, we decided to analyze oligodendrogenesis and neurogenesis separately. In the postnatal forebrain, SOX2 is needed for the expansion of OPCs during developmental and induced myelination and its levels increase at the onset of differentiation and regulate the production and maturation of myelinating oligodendrocytes (Zhang et al., 2018). Reduced SOX2 levels in a p27-null background did not restore the increased production of oligodendrocytes; on the contrary, we detected more newly-generated oligodendrocytes, as reflected in higher numbers of EdU^+^-LRCs in the CC of the compound mutants injected with the nucleoside four weeks before their analysis (**Figure 6J**). Our data indicated that p27 regulates the cycling of OPCs, but the timing of cell cycle exit for differentiation into oligodendrocytes is determined by the level of SOX2. In fetal NPCs SOX2 typically suppresses premature neurogenesis, likely by counteracting the activity of proneural bHLH TFs (Graham et al, 2003; Bylund et al, 2003). In the adult brain, SOX2 also maintains proliferative NPCs undifferentiated and if SOX2 levels are reduced below 50%, the NPC pool is depleted, resulting in impaired neurogenesis (Ferri et al, 2004). In contrast to the oligodendrogenic lineage, SOX2 levels are drastically downregulated at the neuroblast stage, concomitant with a dramatic increase in p27, as shown before. Compound p27KO;*Sox2*^*+/-*^ mice still displayed an increased number of DCX^+^ neuroblasts (**Figure 6K**) but, interestingly, a dosage reduction in SOX2 levels restored p27KO increased numbers of newly-generated OB neurons to WT levels (**Figure 6L**). The data reveal a need for p27-dependent repression of stemness TF SOX2 for neuroblasts to abandon their last cell cycle before they differentiate into OB neurons and suggest that cell cycle withdrawal for terminal neuronal differentiation requires the repression of a stemness-related undifferentiation program. The data together indicate that the regulation of SOX2 dosage by the level of p27 in neuroblasts and OPCs determines cell cycle withdrawal timing and the balance between progenitor cell population expansion and production of specialized cell types.

## Discussion

Arrest of the cell cycle is an obligatory step in NPC differentiation, one which has to be properly coordinated with a switch in gene expression from an undifferentiated to a committed cell-specific profile (Hardwick et al, 2015). We have approached this issue with a combined analysis of the cell state-defining transcriptome and the cell capacity-defining epigenome at the cell transition between mitogen-driven proliferation and mitogen-withdrawal differentiation in the presence and in the absence of the cell cycle inhibitor p27. We provide evidence that increased levels of p27 in response to lack of mitogenic stimulation limit CDK2 activity, restricting cell cycle re-entry and, at the same time, p27 repression of *Sox2* expression acts as a timer for cell cycle arrest for differentiation. Our data indicate that p27 acts as a cell cycle break for NPCs and that *Sox2* upregulation in the absence of p27 contributes to the exuberant neurogenic phenotype of *Cdkn1b* null mice by delaying cell cycle exit of committed neuroblasts. During oligodendrogenesis, precise coordination of p27 and SOX2 levels plays a role in the balance between OPC expansion and oligodendrocyte production. Our results explain how cell cycle exit and the onset of differentiation can be coordinated in the adult SEZ through a mechanism based on a dual p27 action on CDK inhibition and gene expression.

The CKI p27 appears to restrict cell cycling in many cell lineages, as *Cdkn1b* homozygous mutant mice are 30% larger than normal due to hyperplasia in most organs (Fero et al, 1996; Kiyokawa et al, 1996; Nakayama et al, 1996). In line with higher NPC activity in the adult brain, we also found increased numbers of neurons and oligodendrocytes. The apparent discrepancy from a previous study, which described higher NPC activity, but reduced proportions of neuroblasts (Doetsch et al, 2002), most likely reflects actions on apoptosis of the C-terminal domain of p27 (Besson et al, 2008), retained in the specific p27KO (Kiyokawa et al, 1996) analyzed in that report. An increase in p27 levels, as a requirement for cell cycle exit, has been specifically described for several cell types in the CNS (Casaccia-Bonnefil et al, 1997; Gao et al, 1997; Durand et al, 1998). Clonal analyses further indicated that isolated p27-deficient progenitor cells divide one or two times more than wild-type cells before exiting the cell cycle, indicating that other cell cycle components might act as redundant components of the timer or that other cell cycle-independent mechanisms were involved in inducing arrest in a timely fashion (Hardwick et al, 2015; Raff, 2007). However, the underlying mechanism was not fully understood. Analysis at the single-cell level with a CDK2 sensor has allowed us to determine that p27 levels determine the residual CDK2 activity at exit from mitosis and, thereby, the duration of G1 or the propensity of the cell to re-enter a new cell cycle. This could explain why p27-deficient cells can undergo one or two more rounds of cell division in the absence of mitogenic stimulation, resulting in at least a doubling of the resultant population. However, we find that the timing of cell cycle exit for differentiation also requires the regulation of SOX2 levels.

SOX2 belongs to the SOXB1 subfamily, which also includes SOX1 and SOX3 (Reiprich & Wegner, 2015; Sarkar & Hochedlinger, 2013). Although SOX2 is widely known as a pluripotency-sustaining transcription factor in ES cells, it plays a very prominent role in neurogenesis, from specification of the neuroectodermal lineage to maintenance of neural competence (Zhang & Cui, 2014) in a highly dose-dependent manner (Hutton and Pevny, 2011; Baser et al, 2019). Reduction of SOX2 levels limits self-renewal and multipotentiality and promotes NPC cell cycle exit and premature differentiation. In contrast, overexpression of SOX2 not only inhibits the differentiation of multipotential NPCs into neurons and glial cells during fetal development (Bylund et al, 2003; Graham et al, 2003), but can also convert cells of other lineages into NSCs (Ring et al, 2012). Reported dosage effects of SOX2 indicate that its levels are finely regulated (Ferri et al, 2004; Cavallaro et al, 2008; Marques-Torrejon et al, 2013; Taranova et al, 2006); however, its transcriptional and post-transcriptional regulation are still far from understood. Our study and previous ones (Ferri et al, 2004) have demonstrated that reduction of SOX2 levels of as much as 50% does not affect adult neurogenesis in the mouse in a wild-type background, although humans may be more sensitive (Kelberman et al, 2006); however, we have shown that absence of p27 results in increased levels of SOX2 with a clear impact on the neurogenic and oligodendrogenic outputs. A direct role of SOXB1 proteins in cell cycle progression has never been demonstrated, although actions of these regulators in sustaining the undifferentiated state appear linked to a cycling state (Bylund et al, 2003; Graham et al, 2003). We have been able to evaluate the effect of an increased dose of SOX2 and our data also indicate that SOX2 does not play a direct role in cell cycling, but it has to be silenced in neuroblasts to allow their cell cycle exit for differentiation. It has been shown that E2F3a and E2F3b play antagonistic roles in the balance between activation (E2F3b through recruitment of RNA polymerase) and repression (E2F3a together with Rb pocket protein p107) of the *Sox2* gene to achieve a correct dosage in proliferating NPCs (Julian et al, 2013). Modulation of *Sox2* expression by cell cycle-related proteins converts *Sox2* into a key gene involved in coordinating cycling and broad developmental potential.

Our data sustain the concept that SOX2 effects are remarkably dosage-dependent and indicate that the cell context is also essential to explain SOX2 actions. The role of SOX2 in oligodendrogenesis has raised debate until systematic genetic loss-of-function studies have revealed that, while it is involved in the activation of OPCs during demyelinating lesions both in the brain and spinal cord, it is only required for OPC population expansion during developmental myelination in the forebrain (Hoffman et al, 2014; Zhang et al, 2018). SOX2 is subsequently required in a subpopulation of oligodendrocytes during developmental myelination (Zhang et al, 2018), just opposite to the down-regulation in neuroblasts. CKI p27 regulates OPC cycling (Casaccia-Bonnefil et al., 1997; 1999; Gao et al, 1997; Durand et al, 1998; Larocque et al, 2005), including the adult SEZ OPCs as shown here, and it increases in oligodendrocytes where it can enhance expression of the myelin basic protein gene (Miskimins et al, 2002). Our data indicate that moderate levels of p27 act as a cell cycle pacer in OPCs and as a regulator of SOX2 levels. In contrast to neurogenesis, both p27 and SOX2 are required for oligodendrocyte generation and reduction of SOX2 in the absence of p27 retards cell cycle exit of OPCs for differentiation.

Some target genes of SOX2 have been identified in NPCs, including *Egfr*, TLX (*Nr2e1*), and *NeuroD1* (Hu et al, 2010; Kuwabara et al, 2009; Shimozaki et al, 2012). Here we report that SOX2 can directly bind regulatory regions of the *Ascl1* and *Olig2* genes and that their expression directly correlates with SOX2 levels. All these genes are known to regulate the undifferentiated state of NPCs and/or their activity. In addition, some SOX2 target genes encode secreted proteins with direct paracrine and autocrine effects on NSC/NPCs (Favaro et al, 2009) consistent with the idea that SOX2 is a master regulator of expression programs within these cells. Genome-wide binding profile analysis of SOX2 and SOX3 in ES cells, both naïve and specified to NPCs, and in cells differentiated to neurons has revealed an ordered, sequential binding to enhancers associated with a common set of neural genes needed throughout the neurogenic process (Bergsland et al, 2011). Although SOX2 and SOX3 are thought to act redundantly in many contexts, this latter report suggested that SOX2 acts as a pioneer factor that binds neural enhancers for future activation by SOX3, establishing neural competence. Among the silent genes that are pre-bound by SOX2 in ES cells that are later occupied and activated by SOX3 during neural lineage development are *Ascl1* and *Olig2* (Bergsland et al, 2011). Genome-wide analysis of target genes and functional analyses have indicated that ASCL1 plays important roles in proliferating NPCs, promoting the activation of distinct sets of genes during both proliferation and differentiation of NPCs (Castro et al, 2011). Despite its role in neuronal differentiation, ASCL1 can antagonize differentiation programs, i.e. by activating Id1, an inhibitor of bHLH activity through sequestration of E-proteins (Ruzinova & Benezra, 2003), or the co-repressor of proneural proteins CBFA2T2/MTGR1 (Aaker et al, 2010) and activate the expression of cell cycle regulators, possibly in cooperation with MYC (Castro et al, 2011). This duality suggests the existence of mechanisms underlying diversification of ASCL1 actions, which may include association with different partners, post-translational modifications or oscillations in its levels, among others (Castro et al, 2011).

Reprogramming can be used to convert healthy cells in a tissue into a different cell type needed for repair, offering an exciting opportunity for regenerative medicine without the need for transplantation. In the murine CNS, direct reprogramming of endogenous glial cells into viable neurons has been achieved by viral delivery of specific transcription factors such as SOX2, ASCL1 or NEUROD1 (Li & Chen, 2016). SOX2 reprograms somatic cells into pluripotent stem cells and NSCs *in vitro* (Ring et al, 2012) and is sufficient for direct conversion of quiescent mature astrocytes to proliferative ASCL1^+^ NPCs which in turn give rise to DCX^+^ neuroblasts (Niu et al, 2013). Interestingly, upon excitotoxic injury or stroke, striatal astrocytes can re-express ASCL1 and generate neuroblasts (Magnusson et al, 2014; Nato et al, 2015), suggesting the existence of a latent neurogenic program in differentiated astrocytes outside neurogenic niches that can be awakened by the modification of their microenvironment. SOX2-driven reprogramming of astrocytes, therefore, transits through intermediate NPC states before the adoption of a mature neuron fate. *In vivo* conversion and SOX2 actions need, therefore, to be understood at the molecular level. Critically, we have shown here that p27 negatively regulates the transcription of *Sox2* during the course of differentiation into neurons and oligodendrocytes in the adult SEZ independently of its known action on the cell cycle. It may, therefore, be possible to modulate these interactions with small molecules to drive proliferation and/or differentiation in clinical situations to aid repair of damage or restrict tumour growth in the CNS.

## Materials and Methods

### Animals and in vivo manipulations

*Cdkn1b* mice (p27KO) (Fero et al, 1996) were obtained from Jackson Laboratory and maintained on a C57BL/6J background. *Cdkn1b*^*-/-*^*;Sox2*^*+/-*^ mice (D’Amour & Gage, 2003), their control littermates (*Cdkn1b*^*+/+*^*;Sox2*^*+/+*^, *Cdkn1b*^*-/-*^*;Sox2*^*+/+*^) and Rosa26R-loxP-STOP-loxP-*Sox2*-GFP mice (for *Sox2* overexpression) (Lu et al, 2010) were maintained on mixed backgrounds. All comparisons were made among mice derived from the same sets of crosses and they shared the same genetic background. Animals were genotyped by PCR analysis of DNA extracted from mouse ear-punch using specific primers (*Wild-type*-F: GATGGACGCCAGACAAGC; *Wild-type*-R: CTCCTGCCATTCGTATCTGC; *Cdkn1b*-F: CTTGGGTGGAGAGGCTATTC; *Cdkn1b*-R: AGGTGAGATGACAGGAGAT) or done by Transnetyx. Housing and experiments were carried out following protocols approved by the Ethics Committee of the Universidad de Valencia (CEEA: A1403250608615) (Spain) and by the Animal Ethics Committee and the UK Home Office (PPL 70/8560) at the Francis Crick Institute (London, UK).

### Tissue preparation and immunohistochemistry

BrdU administration regimes have been previously detailed (Ferron et al, 2007). Mice were injected intraperitoneally (i.p.) with 50 mg of BrdU (Sigma, B5002) *per* kg of body weight every 2 h for 12 consecutive hours (7 injections in total) and euthanized either immediately or 28 days later or with just one injection one hour before euthanasia. EdU (Life Technologies, E10187) was administered i.p. at 30 mg/kg every 12 h for 3 consecutive days (6 injections in total) and mice were euthanized 28 days after the last injection. Mice were deeply anesthetized and transcardially perfused with 100 ml of 4% paraformaldehyde (PFA) in 0.1 M phosphate buffer pH 7.4 (PB). For ASCL1 immunodetection, less than 40 ml of PFA were infused. Brains were dissected out and vibratome-sectioned at 40 μm. For immunohistochemistry, sections were washed in PBS and blocked at room temperature for 1h in PBS (0.9% NaCl in PB) with 0.2% Triton X-100 supplemented with 10% FBS and then incubated overnight at 4 °C with primary antibodies (1:100 mouse anti-ASCL1, BD 556604; 1:800 rat anti-BrdU, Abcam ab6326; 1:400 chicken anti-DCX, Abcam ab153668; 1:300 goat anti-DCX, Santa Cruz sc-8066; 1:800 chicken anti-GFAP, Millipore AB5541; 1:300 rabbit anti-Ki67, Abcam ab15580; 1:500 rabbit anti-OLIG2, Millipore AB9610; 1:200 rabbit anti-p27, Cell Signaling 3686; 1:700 mouse anti-PSA-NCAM, Millipore MAB5324; 1:600 goat anti-SOX2, R&D AF2018). For BrdU detection, sections were pre-incubated in 2N HCl for 20 min at 37 °C and neutralized in 0.1 M sodium borate (pH 8.5). Detections were performed with 1:800 fluorescent secondary antibodies (AF488 donkey anti-chicken, Jackson ImmunoResearch 703-545-155; AF488 donkey anti-rabbit, Jackson ImmunoResearch 711-547-003; AF647 donkey anti-goat, Molecular Probes A21447; Cy3 donkey anti-chicken, Jackson ImmunoResearch 703-165-155; Cy3 donkey anti-mouse, Jackson ImmunoResearch 715-165-151; Cy3 donkey anti-rabbit, Jackson ImmunoResearch 711-165-152; Cy3 donkey anti-rat, Jackson ImmunoResearch 712-165-153). EdU detection was carried out using the Click-iT™ Plus EdU Alexa Fluor™ 555 Imaging Kit (ThermoFisher, C10638). Nuclei were counterstained with 1 µg/ml of DAPI and sections were mounted with Fluorsave (Calbiochem).

### Neurosphere cultures, differentiation and immunocytochemistry

Methods for NSC culture derived from SEZ, self-renewal assessment, differentiation and for BrdU immunocytochemistry in neurosphere cultures have been previously described in detail (Belenguer et al, 2016; Ferron et al, 2007). For proliferation assays, NSCs were grown in neurosphere control medium containing 20 ng/ml epidermal growth factor (EGF) (Invitrogen, 53003-018) and 10 ng/ml fibroblast growth factor (FGF) (Sigma, F0291). Importantly, for bulk differentiation assays on secondary neurospheres, 40,000 cells/cm^2^ were seeded for 2 days (2 DIV) in Matrigel®-coated coverslips and in control medium with FGF. After these 48 h, mitogens were removed and cells were grown in control medium supplemented with 2% Fetal Bovine Serum (FBS). Differentiating cultures were analyzed 24 h (2+1 DIV), or 5 days (2+5 DIV) after mitogen withdrawal (see **Figure 1A**) depending on the experimental procedure. For immunocytochemistry, cultures were fixed with 2% PFA in PB for 15 min, blocked (PBS with 0.1% Triton X-100 and 10% FBS) and incubated with primary (overnight at 4 °C) and secondary (1h at room temperature) antibodies (1:100 mouse anti-ASCL1, BD 556604; 1:100 mouse anti-ASCL1, Guillemot lab; 1:800 rat anti-BrdU, Abcam ab6326; 1:800 chicken anti-GFAP, Millipore AB5541; 1:300 rabbit anti-Ki67, Abcam ab15580; 1:300 mouse anti-O4, Hybridoma Bank rip; 1:500 rabbit anti-OLIG2, Millipore AB9610; 1:200 rabbit anti-p27, Cell Signaling 3686; 1:200 mouse anti-p27, Cell Signaling 3698; 1:600 goat anti-SOX2, R&D AF2018; 1:300 rabbit anti-ßIII-TUBULIN, Sigma T2200; 1:800 AF488 donkey anti-mouse, Molecular Probes A21202; 1:800 AF488 donkey anti-rabbit, Jackson ImmunoResearch 711-547-003; 1:800 AF647 donkey anti-goat, Molecular Probes A21447; 1:1000 biotin horse anti-mouse, Vector Laboratories BA-2000; 1:2000 Cy3-streptavidin, Jackson ImmunoResearch 016-160-084; 1:800 Cy3 donkey anti-chicken, Jackson ImmunoResearch 703-165-155; 1:800 Cy3 donkey anti-mouse, Jackson ImmunoResearch 715-165-151; 1:800 Cy3 donkey anti-rabbit, Jackson ImmunoResearch 711-165-152; 1:800 Cy3 donkey anti-rat, Jackson ImmunoResearch 712-165-153). DAPI (1 µg/ml) was used to counterstain nuclei. 10 µM EdU was administered for 1 h prior to fixation and detected using the Click-iT™ Plus EdU Alexa Fluor™ 555 Imaging Kit (ThermoFisher Scientific, C10638) following the manufacturer’s instructions. For CDK2 inhibition, 2 DIV differentiated cells were treated with 1 µM CDK1/2 inhibitor III (Millipore, 217714) in control medium supplemented with 2% FBS for 24 h. An appropriate dilution of DMSO was used as vehicle.

### Flow cytometry

Characterization of NSC and NPC populations in the adult SEZ was performed as previously described (Belenguer et al, 2021a,b). After dissection, both SEZs from each mouse were minced and enzymatically digested using the Neural tissue dissociation kit (T) (Miltenyi, 130-093-231) following the instructions of the manufacturer in a gentleMACS Octo Dissociator with heaters (Miltenyi). Digestion was quenched with 100 μg/ml trypsin inhibitor and the digested pieces were mechanically dissociated pipetting up and down 20 times through a plastic Pasteur pipette. Cell suspension was filtered through a 40 μm nylon filter and cells were pelleted (300 x*g*, 10 min) and incubated with the Dead Cell Removal Kit (Miltenyi, cat no. 130-090-101) following the instructions of the manufacturer. Finally, the eluted living fraction was pelleted (300 x*g*, 10 min), resuspended in 100 μl blocking buffer (HBSS without Calcium and Magnesium, 10 mM HEPES, 2 mM EDTA, 0.1% Glucose, 0.5% BSA) and incubated with the specific primary antibodies (1:300 CD24-PerCP-Cy5.5, BD 562360; 1:100 CD31-BUV395, BD 740239; 1:200 CD45-BUV395, BD 565967; 1:20 CD9-Vio770, Miltenyi 130-102-384; 1:20 GLAST-PE, Miltenyi 130-095-821; 1:30 O4-Biotin, Miltenyi 130-095-895; 1:200 Ter119-BUV395, BD 563827; 1:300 AF488 EGF complex, Molecular Probes E13345) and reagents (DAPI, 50 µg/ml) at 4 °C for 30 min. After washing with 1 ml of blocking buffer, labeled samples were centrifuged (300 x*g*, 10 min at 4 °C) and resuspended in 0.5 ml of blocking buffer. Labeled samples were analyzed using a LSR-Fortessa cytometer (Becton Dickinson) with 350, 488, 561 and 640 nm lasers.

### Chromatin immunoprecipitation (ChIP)

ChIP was performed essentially as previously described (Strogantsev et al, 2015). Wild-type neurospheres (3 DIV) isolated from the adult SEZ and 2+1 DIV differentiated cells were cross-linked and chromatin isolated. Chromatin was sheared to an average size of 200-500 bp using a Bioruptor sonicator (Diagenode, UCD-200). Chromatin was pre-cleared with 20 µl of protein G magnetic beads (Dynabeads®, 10003D) for 3 h at 4 °C with rotation. 5 % of chromatin after pre-clearing was used as input. 5-10 µg of antibody (rabbit anti-p27, Santa Cruz sc-528; goat anti-SOX2, R&D AF2018) or non-related antibody (NRA) were added and incubated overnight at 4 °C on a rotation wheel. Neu (C-18) (rabbit anti-NEU, Santa Cruz sc-284) and Rock2 (N-19) (goat anti-ROCK2, Santa Cruz sc-1852) were used as NRA for p27 and SOX2 immunoprecipitations, respectively. Chromatin was precipitated with 10 µl protein G beads for 3 h at 4 °C. Beads were then washed followed by crosslink reversal and protein digestion. Finally, DNA was purified using MinElute PCR purification kit (Qiagen, 28004) following manufacturer’s instructions. ChIP enriched DNA was analyzed by real-time PCR using SYBR-green primers (prAscl1[-906/-784]-F: TAACCCTGAGTGCCTTCCTG; prAscl1[-906/-784]-R: GGAGTCATGAACAGGATAGGTTG;; prAscl1[-523/-432]-F: CTGCGGAGAGAAGAAAGGGG; prAscl1[-523/-432]-R: TCAGGGAAGGGTTTAGGCAG; prOlig2[-637/-546]-F: CTGCAGCAACTGCCACTAAG;; prOlig2[-637/-546]-R: CTGTGACCATTTGTGGTTGC; prOlig2[-147/-58]-F: TTCATTGAGCGGAATTAGCC; prOlig2[-147/-58]-: CGGGAACAATGTGCTTTTC; prSox2[-460/-345]-F: ATGAGCGCAGAAACAATGGCA; prSox2[-460/-345]-R: ACATAAGGGTGGATGGGGCG; SRR2[+4042/+4133]-F: AAGAATTTCCCGGGCTCG; SRR2[+4042/+4133]-R: CCTATGTGTGAGCAAGAACTGTCG;). Pull-downs using NRA were used to control for non-specific enrichments. The comparative Ct method was used to calculate fold enrichment levels relative to NRA after normalizing each one to input.

### Transfection and viral infection of NSCs

For restoration of p27 and SOX2 levels in *Cdkn1b*^*-/-*^ cultures, NSCs were transfected using the Amaxa NSC Nucleofector Kit (Lonza, VPG-1004) as previously described (Belenguer et al, 2016) with either 7.5 μg of pcDNA3.1 and pcDNA3.1-p27-Flag (p27 full-length construct) or 7.5 μg of empty pMSCVpuro (Clontech, 634401) and pSUPER.retro mSox2.3 (*sh*RNA*Sox2* target sequence: 5’-CGAGATAAACATGGCAATCAA-3’), respectively. *Cdkn1b*^*+/+*^ cultures were used as control. Nucleofected cells were plated on differentiative conditions and processed for immunocytochemistry at 2 and 2+1 DIV. For overexpression of *Sox2*, Rosa26R-loxP-STOP-loxP-*Sox2*-GFP NSC cultures were infected with Ad-GFP and Ad-CMV-iCre (Vector Biolabs) following the protocol described in (Hawley et al, 2010). These cells carry the endogenous *Sox2* alleles plus a conditional allele of the *Rosa26* locus containing a Sox2-IRES-GFP cDNA and a synthetic cytomegalovirus early enhancer/chicken beta actin (CAG) promoter upstream of *Sox2* to increase the expression. Shortly, 30,000 individual NSCs were infected with a multiplicity of infection (MOI) of 500 in complete medium. After 24 h, viruses were washed and cells were plated in fresh medium and left to grow for 5 more days. After passage, cells were plated on differentiative conditions and processed for immunocytochemistry at 2 and 2+1 DIV. Cre-infected cultures are referenced as *Sox2*^*OE*^ while control GFP-infected are noted as *Sox2*^*+/+*^ in those experiments. The 30% brightest cells in both conditions were considered as SOX2^high^. In order to study CDK2 activity, *Cdkn1b*^*+/+*^ and *Cdkn1b*^*-/-*^ cultures were infected with the lentiviral particles carrying the CSII-EF-DHB-mVenus construct (Spencer et al, 2013) as previously described (Porlan et al, 2014). Infected neurospheres were cultured for 5 days in fresh complete medium, split and differentiated according to the experimental procedure.

### Cloning and Luciferase assay

Murine *Sox2* proximal promoter (positions -1907 to +6 relative to the transcription initiation site) (Wiebe et al, 2000) was retrieved by PCR from C57B6/J genomic DNA with specific primers containing restriction-site target sequences (*prSox2*[-1907/+6]-luc-F: AAAAAACTCGAGAAACTTAAGGAGAACCTGGGG; prSox2[-1907/+6]-luc-R: CCACCAAAGCTTAACAAGTTAATAGACAACCATCCA). Purified and digested DNA fragment was cloned into a pGL3-Enhancer vector (Promega) and the correct sequence was checked by Sanger sequencing. *Cdkn1b*^*+/+*^ and *Cdkn1b*^*-/-*^ cells were nucleofected (see ***Transfection and viral infection of NSCs***) with 7.5 μg pCDNA3.1 or pCDNA3.1-p27-Flag, 0.5 μg of pMAX-GFP and 2 μg of the corresponding reporter construct (*prSox2*[-1907/+6]-luc; *SRR2*[+3641/+4023]-luc, kindly provided by Dr. Okuda (Tomioka et al, 2002), driving the expression of the firefly luciferase and *Renilla* luciferase plasmid in a 1:20 ratio. After electroporation, NSCs were plated on either proliferative or differentiative conditions. Cells were lysed after 72 h (2+1 DIV for differentiations) using the Dual Luciferase Reporter kit (Promega, E1960) and luciferase activity was measured using a Victor3 (Perkin Elmer) reader. Ratio of firefly to *Renilla* luciferase was calculated and represented as arbitrary units (a.u.).

### Imaging and bioimage analysis

Images were acquired with an Olympus FV10i confocal microscope (Olympus). For *in vitro* experiments, laser settings were first established on wild-type/control samples at the first time point (e.g. 2 DIV) and kept throughout the whole experiment. Random fields were imaged at the focal plane showing both the highest number of cells on focus and the most intense signal. 2+5 DIV differentiation samples were imaged using a Nikon ECLIPSE Ni-U microscope (Nikon) with a Zyla 4.2 sCMOS camera (Andor). Bioimage analysis was performed using the Fiji open-source software package (Schindelin et al, 2012). ImageJ Macro Language scripts were developed in order to automatically perform an unbiased analysis of some of the cellular assays. On one hand, for the analysis of EdU pulse-chase experiments, as well as the signal measurement of other nuclear proteins labeled by means of immunostaining (Ki67, SOX2, OLIG2, ASLC1), the ‘Cell proliferationHCS’ tool was used (Carrillo-Barberà et al, 2019). A signal threshold was set for each marker to classify cells as positive. A minimum of 500 cells per culture were scored in all conditions. On the other hand, CDK2 activity was also analyzed by means of a bioimage analysis workflow (see **Figure 2D**) that obtains a binary mask for each cell in both DAPI and CSII-EF-DHB-mVenus images. After that, nuclear particles of non-infected cells (mVenus^-^) are discarded. The remaining particles are submitted to a Voronoi partition in order to separate touching or clumped cells. To obtain a mask containing just the mVenus-positive (mVenus^+^) cytoplasm, nuclear masks are erased from the mVenus binary mask by means of the Boolean operator XOR. Cytoplasmic particles smaller than their corresponding nucleus are discarded. Finally, nuclear and cytoplasmic binary masks are redirected to the original grayscale image to retrieve the mVenus signal (mean grey value and integrated density). Cells with an mVenus cytoplasm/nucleus ratio (mean gray value) lower than 0.65 were classified as G0/G1 cells, whereas those without mVenus^+^ cytoplasm mask were directly scored as G0/G1. Then, to measure the residual CDK2 activity during this phase, the integrated intensity of the mVenus positive cytoplasms of cells in G0/G1 was scored. A minimum of 250 cells in G0/G1 per culture were scored in all conditions. Full script can be obtained from GitHub (https://github.com/paucabar/DHB-Venus). For *in vivo* experiments, laser settings were first established on wild-type tissue and similar regions of interest (ROI) were acquired in an Olympus FV10i confocal microscope. Maximal projection images were generated and the mean grey intensities of nuclear markers (ASCL1, OLIG2, SOX2) were measured with ImageJ/Fiji software. Intensities were represented as frequency histograms normalized to the maximum count in each comparison.

### RNA- and ATAC-seq

3 DIV proliferating neurospheres and NPs at the onset of differentiation (2+1 DIV) were harvested for RNA and chromatin extraction. For ATAC sequencing 50,000 cells were centrifuged (500x*g*, 5 min, 4 °C), washed with ice-cold PBS and lysed in cold lysis buffer (10 mM Tris-HCl pH 7.4, 10 mM NaCl, 3 mM MgCl2, 0.1% IGEPAL CA-630). Nuclei were pelleted (500x*g*, 10 min, 4 °C) and tagmented with the Nextera DNA Library Prep Kit (Illumina, FC-121-1030) for 1 h at 37 °C. After transposition, DNA was purified with the MinElute PCR purification kit (Qiagen, 28004) and amplified and barcoded with the NEBNext High-Fidelity 2X PCR master mix (New England Labs, M0541) following manufacturer’s instructions. The amplified library was then purified with XP AMPure Beads (Beckman Coulter, A63880). Library fragments were analyzed and quantified with a 2100 Bioanalyzer (Agilent Technologies). Libraries were sequenced on an Hiseq4000 (Illumina) to achieve an average of 25 million reads per sample in the Advanced Sequencing Facility (ASF) at the Francis Crick Institute. For bulk RNA sequencing, samples were lysed with the QIAzol Lysis Reagent (Qiagen, 79306) and RNA was extracted with the miRNeasy micro kit (Qiagen, 217084) according to the manufacturer’s protocol. RNA quality was assessed using the Agilent RNA 6000 Pico Kit (Agilent Technologies). cDNA was generated using Ovation RNA-seq System V2 (Tecan, 7102-A01), libraries were constructed using Ovation Ultralow System V2 (Tecan, 0344NB-A01) according to the manufacturer’s instructions. Libraries were quantified using the TapeStation (Agilent), pooled in equimolar proportions, and sequenced on an Hiseq4000 (Illumina) to achieve an average of 25 million reads per sample.

### Bioinformatic analyses

#### ATAC sequencing

Paired-end sequencing files were analyzed with FastQC, summarized with MultiQC and visually inspected for major quality issues. We used Cutadapt to clip the Nextera 3’ R1 and R2 adapters and then Trimmomatic to trim low-quality portions and filter reads shorter than 20 bases. A second round of quality control with FastQC and MultiQC was performed after the initial filtering steps to make sure the outputs were in line with the expected results. High quality reads were then mapped to the mouse genome with Bowtie2, using the *Mus musculus* GRCm38 (mm10) genome as reference and allowing mate dovetailing. After mapping, duplicates were marked on BAM files with MarkDuplicates from Picard tools, insert sizes were calculated with the CollectInsertSizeMetrics tool from SAMtools, and then peak calling was performed with MACS2, choosing narrowpeaks, no-model and 0.05 FDR as options. After peak calling, the resulting narrow-peak BED files were manually loaded into the IGV genome browser alongside their corresponding ATAC-seq coverage tracks generated with the bamCoverage function from DeepTools for visual inspection. Biological replicates of each sample were merged and used to generate average metagene plots and per-gene accessibility heatmaps around transcription start site (TSS) genomic coordinates of mouse protein-coding genes with ngsplot. MACS2 narrow peak files and the corresponding BAM alignment files they were derived from were used as input for differential peak calling with DiffBind. An FDR threshold of 0.05 was chosen to decide whether a peak was differentially detected between comparisons. Peaks that were significantly more or less concentrated in one sample compared to its control sample were considered Open and Closed, respectively. In order to associate both differential (Open and Closed) and non-differential peaks to their genomic contexts, we used ChIPseeker with the Gencode M18 mouse GTF annotation. Upstream and downstream regions of 3 Kb were allowed, as well as a flanking gene distance option of 5 Kb. To simplify the analysis and more directly associate changes in ATAC signal to regulatory events in downstream genes, we focused on peaks associated with 5’ and promoter regions, as determined by the ChIPseeker annotation, for downstream analysis. In the cases where there was more than one promoter peak associated with the same gene, we kept either the one that was closest to the TSS, or the largest overlapping one. Differential promoter peak coordinates were used as input for the DNA motif analysis tool HOMER, with the findMotifsGenome.pl Perl script and the -size given option, which instructs the algorithm to look for motifs in all the peak, not just in the region around the center of the peak.

#### RNA sequencing

Raw sequencing files were aligned to the Ensembl mouse transcriptome using STAR, and then RSEM was used to count raw reads per gene. Genes with less than 1 count per million (CPM) in at least 4 of the samples were not considered for further analysis. Genomic coverage tracks were generated from BAM alignment files with the bamCoverage function from DeepTools, choosing RPKM normalization as a scaling option. Coverage tracks were loaded into the genome browser IGV for visual inspection. Differential expression analysis between samples was done with DESeq2, and an FDR threshold of 0.05 was used to discriminate between UP- or DOWN_regulated genes. Similarly to ATAC-seq samples, the biological replicates of each RNA-seq sample were merged and used as input for ngsplot to generate average metagene plots and per-gene mRNA amount heatmaps around transcription start site (TSS) genomic coordinates of protein-coding genes.

#### Integration between ATAC-seq and RNA-seq

The tabular outputs of DiffBind (ATAC-seq) and DESeq2 (RNA-seq) were loaded in R for further processing. We combined both matrices and labeled every gene with its corresponding differential accessibility (DA) and differential expression (DE) analysis results. This way genes were tagged as DA *Closed, Open or Unchanged*, and as DE *UP, DOWN* or *Unchanged*. Out of the nine possible combinations of differential traits, we focused for downstream functional analysis on only genes with significant differences in both modalities simultaneously: *Closed_UP, Closed_DOWN, Open_UP* and *Open_DOWN*. Functional enrichment analysis of differential genes was carried out using a variety of tools, including STRING, Mousemine, GOrilla, REVIGO, Panther, Enrichr, Appyter and the ClusterProfiler R package. The AnimalTF and UNIPROT databases were used to extract information about mouse TFs and TF families. The Epigenetic Landscape In Silico deletion Analysis (LISA) tool was used to discover associations between our target gene lists and regulatory TFs and chromatin factors based on the Cistrome DNase and ChIP-seq database (CistromeDB). Density scatterplots, Venn diagrams and violin plots were generated in R with custom code.

### Statistical analysis

All statistical tests were performed using the GraphPad Prism Software, version 5.00 for Windows (http://www.graphpad.com). Analyses of significant differences between means were carried out using the unpaired or paired two-tailed Student t-test or one-way ANOVA with Tukey post-hoc test when appropriate. When comparisons were carried out with relative values (normalized values and percentages), data were first normalized by using a log or arcsin transformation, respectively. Data are always presented as the mean ± standard error of the mean (s.e.m). The number of experiments carried out with independent cultures/animals (n) is either shown as dots in the graphs or listed in the Figure Legends. Statistical significance in the violin-plot was assessed by the Wilcoxon signed-rank test as implemented in the ggpubr R package (\**P* < 0.05, \*\**P* < 0.01, \*\*\**P* < 0.001, and \*\*\*\**P* < 0.0001).

## Acknowledgements

We thank M. J. Palop for help with the mouse colonies and acknowledge the support of the Servicio Central de Soporte a la Investigación Experimental (SCSIE-UVEG). We would also like to thank Dr. F. Guillemot (Francis Crick Institute, London) for providing the ASCL1 antibody and Drs. Jesus Gil (Imperial College, London), Anxo Vidal (Centre for Research in Molecular Medicine and Chronic Diseases, Santiago de Compostela), Akihiko Okuda (Research Center for Genomic Medicine, Saitama), and Tobias Meyer (Stanford University Medical Center, Stanford) for generously providing DNA constructs. This work was supported by grants PID2020-117937GB-I00, RED2018-102723-T, CB06/05/0086 (CIBERNED), and RD16/0011/0017 (RETIC Tercel) from Ministerio de Ciencia e Innovación and Prometeo 2021/028 from Generalitat Valenciana to I.F. and core funding from the Francis Crick Institute (FC001107, from Cancer Research UK, the UK Medical Research Council, and Wellcome) and an NIH grant (R01-EB016629) to R.L-B. P.C-B. was recipient of a MICINN’s FPI predoctoral contract and A.D-M. and G. B. were recipients of FPU predoctoral contracts from Ministerio de Educación, Cultura y Deporte (MECD). A.D-M.’s stay at R.L-B.’s laboratory was supported by a MECD’s grant included in the FPU program.

## Author contributions

Conceptualization, A.D-M., JM.M-R., O.B., SR.-F., R.L.-B., I.F.; Methodology, A.D.-M., JM.M-R., V.M.-A., SR.-F., G.B..; Formal Analysis, A.J-P., P.C.-B.; Investigation, A.D.-M., JM.M-R., V.M.-A., SR.-F., P.C.-B., G.B., A.P.-V., E.P., M.K.; Resources, V.M.-A., R.L.-B.; Data Curation, A.J-P.; Writing – Original Draft, I.F., JM.M.-R., A.D.-M.; Writing – Review & Editing, A.D-M., JM.M-R., SR.-F., A.J.-P., V.M.-A., I.F. R.L.-B.; Visualization, A.D-M., JM.M-R., A.J.-P.; Supervision, I.F.; Project Administration, I.F.; Funding Acquisition: I.F., R.L.-B.

## Competing financial interest statement

The authors declare no competing financial interests.

## Figure and Table Legends

**Table S1**. LISA results for the comparison of Closed_DOWN vs Closed_UP genes in WT cells.

**Table S2**. LISA results for the comparison of Closed (UP+DOWN) in WT vs Closed (UP+DOWN) genes in p27KO cells.

## Notes

### Competing Interest Statement

The authors have declared no competing interest.

